# Insects as phyllosphere microbiome engineers: effects of aphids on a plant pathogen

**DOI:** 10.1101/797738

**Authors:** Melanie R. Smee, Imperio Real-Ramirez, Tory A. Hendry

**Affiliations:** Department of Microbiology, Cornell University, Ithaca, NY 14853, USA

**Keywords:** Aphids, environmental bacteria, honeydew, plant-microbe-insect interactions, plant pathogens, *Pseudomonas syringae*

## Abstract

Insect herbivores are common in the phyllosphere, the above-ground parts of plants, and encounter diverse plant-associated bacteria there, yet how these organisms interact remains largely unknown. Strains of the bacterium *Pseudomonas syringae* grow well epiphytically and have been shown to grow within and kill hemipteran insects like the pea aphid, *Acyrthosiphon pisum*. Aphids are hypothesized to be an alternative host for these epiphytic bacteria but it is unclear if aphids provide fitness benefits to these bacterial pathogens. To determine if epiphytic bacteria could be adapted for infecting aphids, we characterized 21 strains of *P. syringae* for epiphytic ability and virulence to pea aphids and found that the two traits were positively correlated. For a subset of strains, we tested if the bacteria derived a fitness benefit from the presence of aphids. Some strains benefited significantly, with up to 18.9% higher population densities when aphids were present, and lower starting population density was predictive of higher benefit from aphid presence. However, further investigation found that honeydew, the sugary waste product of aphids, and not growth in aphids, increased *P. syringae* growth on leaves. This suggests that aphids may be important microbiome engineers in the phyllosphere, but evolutionarily dead-ends for epiphytic bacteria.

## 1 Introduction

The phyllosphere, consisting of all the aerial parts of plants, is a complex and dynamic environment (Lindow and Brandl 2003; Vorholt 2012; Farré-Armengol et al. 2016; Laforest-Lapointe et al. 2017; Saleem et al. 2017). Estimated at having approximately double the global surface area of the land, the upper and lower leaf surfaces are an important, yet heterogeneous, environment (Vorholt 2012). Leaf surfaces provide the stage for a multitude of interactions between plants, animals and microbes. Work on these interactions shows co-evolutionary dynamics between microbes and hosts (Koskella and Brockhurst 2014; Flórez et al. 2017), but many such systems, especially tripartite interactions, are understudied despite their potential importance in diverse systems. Some species of plant-associated bacteria form epiphytic populations on leaf surfaces, with densities averaging 10^5^ - 10^7^ cells/cm^2^ (Beattie and Lindow 1995; Andrews and Harris 2000; Hirano and Upper 2000; Monier and Lindow 2004). For phytopathogens, such high densities can lead to an increased chance of infection (Hirano and Upper 2000), and epiphytic bacteria likely influence plants in other ways as well (Hornschuh et al. 2002; Abanda-Nkpwatt et al. 2006). At a local scale, communities of epiphytic bacteria are often dominated by a few bacterial taxa (Bulgarelli et al. 2013), yet at a broader scale, even on a single plant, bacterial communities can be diverse and variable (Kembel et al. 2014; Meaden et al. 2016). Hence, it can be difficult to explain observed bacterial distributions.

Attempts to uncover what generates and maintains microbial diversity in the phyllosphere have explored various abiotic and biotic factors. Frequently changing, and often extreme, temperatures, humidity levels, radiation, and limited nutrients all combine to produce a challenging habitat (Hirano and Upper 2000), as well as competition from other microbial species (Wilson and Lindow 1994), and even interactions with bacteriophages (Morella et al. 2018). Animals can also have an impact on the distribution of microbes in the phyllosphere, such as vectoring of pathogens on crops (Perilla-Henao and Casteel 2016). Although phyllosphere microbes are known to impact animal hosts (Leroy et al. 2011*a*; Kupferschmied et al. 2013; Mason et al. 2014; Farré-Armengol and Junker 2019), few studies have considered the potential ecological or evolutionary implications of interactions with animals for the microbe (Biere and Bennett 2013). To further complicate matters, tripartite interactions in the phyllosphere often involve plant-mediated traits, which may be difficult to discern from direct interactions (Stout et al. 2005; Humphrey et al. 2014). For instance, herbivores can affect the growth of microorganisms on forest trees (Stadler and Müller 2000), and in particular, leaves damaged by lepidopteran larvae host a greater diversity of bacteria, or a change in bacterial community composition (Müller et al. 2003). However, it is not known if this effect is plant-mediated.

The pea aphid, *Acyrthosiphon pisum*, is a major agricultural pest and acts as both potential host and vector for some strains of *Pseudomonas syringae*, a ubiquitous plant-associated bacterium and phytopathogen (Stavrinides et al. 2009; Smee et al. 2017). Along with a common host plant, *Vicia faba*, they are a tractable system for exploring questions about tripartite interactions in the phyllosphere. *P. syringae* is a diverse species complex in the Gammaproteobacteria, which can differ greatly in their phenotypes despite being very phylogenetically close (Baltrus et al. 2012). Strains vary dramatically in their ability to colonize plants epiphytically, but also in their virulence to both their plant host and to *A. pisum*. Some strains are able to cause the death of up to 83% of aphids in a 72 hour period (Smee et al. 2017). The variation in virulence to *A. pisum* across 12 strains of *P. syringae* was found to be correlated with epiphytic ability of the strains (Smee et al. 2017). Yet it is unclear why this correlation exists, and why virulence to hemipteran insects would be adaptive for epiphytic bacteria. Several strains of *P. syringae* replicate to high densities (10^6^ to 10^8^ CFUs per aphid) within aphid hosts and are also found in the honeydew of aphids infected with at least one pathovar, *P. syringae* pv. *syringae* B728a (from here on referred to as PsyB728a) and deposited onto plants (Stavrinides et al. 2009; and unpublished data). Perhaps the ability of bacteria to grow on leaf surfaces may better allow them to infect transient insects visiting plants, but in that case has high virulence to insects been selected for in epiphytic strains? Do such strains experience a trade-off between virulence and dispersal, where growing inside an aphid for longer could allow for more dispersal? To address these evolutionary questions, we must first understand if the fitness of epiphytic bacteria is indeed influenced by interactions with insect hosts.

Firstly, we undertook a robust test of the hypothesis that epiphytic ability and virulence to aphids are correlated, using a total of 21 *P. syringae* strains. Secondly, we determined whether this correlation may be adaptive by testing if *P. syringae* benefits from using aphids as hosts, by comparing epiphytic bacterial densities on plants with and without aphids. We lastly explored what traits may influence this benefit, in the absence of plant-mediated effects, by separating fitness benefits from growth inside insects and the benefits derived from changes that insects create in the phyllosphere, such as deposition of aphid honeydew, a sugar-rich waste product.

## 2 Materials and Methods

### 2.1 Insect and bacterial cultures

The pea aphid, *A. pisum*, reproduces parthenogenetically under summer conditions, so lab populations were kept on fava bean (*V. faba*) plants at 20°C and a light:dark 16:8 hour cycle. We used *A. pisum* clone CWR09/18 collected by Angela Douglas in Freeville, NY in 2009, which does not harbor any bacterial symbionts aside from the obligate symbiont *Buchnera aphidicola* (Macdonald et al. 2012). Aphids were age-controlled on *V. faba* for all experiments so that all individuals were 5-6 days old at the start of each assay.

All strains of *P. syringae* used in the current study displayed resistance to the antibiotic rifampicin, apart from one, pv*. syringae* B301D. Therefore, all bacterial cultures were grown and kept on King’s B (KB) media with added rifampicin (50 ng/μl) apart from pv. *syringae* B301D which did not have rifampicin added. Culture plates were kept incubated at 27°C and overnight cultures were grown in an incubator shaker set at 28°C and 220 rpm.

### 2.2 In vitro pathogenicity assays

Oral infection assays were used to test the virulence of a total of 21 strains of *P. syringae* (Table 1). These strains represent the phylogenetic breadth of plant pathogenic strains commonly included within the species complex *P. syringae* (phylogroups 1, 2, 3 and 5 are included), with the highest representation within phylogroup 2, which has previously been found to include highly epiphytic and insect virulent strains (Smee et al. 2017). We note that some strains are considered distinct species in some classifications (Sarkar and Guttman 2004; Bull et al. 2011; Borschinger et al. 2016). Additionally, some of the included strains are environmental isolates from water sources or are thought to be non-pathogenic to plants. All isolates were provided by David Baltrus and references for origins of strains are given in Table 1.

**Table 1.** A summary of *P. syringae* strains used in the current study. MLST phylogroup designation and original source are given, with GenBank accession numbers for genome sequences used in phylogenetic tree construction (Borschinger et al. 2016). Those strains for which data are already reported by Smee *et al.* (2017) are highlighted in grey.

| Name | Pathovar | Strain | Group | Source | Genbank accession |
| --- | --- | --- | --- | --- | --- |
| PtoDC3000 | *tomato* | DC3000<sup>*1</sup> | 1 | *Solanum lycopersicum* | AE016853 |
| Pja301072 | *japonica* | MAFF 301072 PT<sup>*2</sup> | 2 | *Hordeum vulgare* | AEAH 00000000 |
| Ptt50252 | *aptata* | DSM50252<sup>*3</sup> | 2 | *Beta vulgaris* | AEAN 00000000 |
| Ppi1704B | *pisi* | 1704B<sup>*3</sup> | 2 | *Pisum sativum* | AEAI 00000000 |
| PsyCC1543 | *syringae* | CC1543<sup>*4</sup> | 2 | Fresh lake water | AVEJ 00000000 |
| PsyCC1458 | *syringae* | CC1458<sup>*4</sup> | 2 | *Dodecantheon pulchellum* | AVEN 00000000 |
| PsyCC440 | *syringae* | CC440<sup>*5</sup> | 2 | *Cucumis melo* | AVEC 00000000 |
| PsyCC457 | *syringae* | CC457<sup>*5</sup> | 2 | *Cucumis melo* | AVEB 00000000 |
| PsyB728a | *syringae* | B728a<sup>*6</sup> | 2 | *Phaseolus vulgaris* | CP000075 |
| PssB48 | *syringae* | B48<sup>*7</sup> | 2 | *Prunus persica* | LGKT 00000000 |
| PsyB301D | *syringae* | B301D | 2 | *Pyrus communis* | CP005969 |
| Psy1212 | *syringae* | 1212 | 2 | *Pisum sativum* | AVCR 00000000 |
| PsyUB303 | *syringae* | UB303<sup>*8</sup> | 2 | Fresh stream water | AVDZ 00000000 |
| PsyUSA011 | *syringae* | USA011<sup>*5</sup> | 2 | Fresh stream water | AVDX 00000000 |
| Pac302273 | *aceris* | MAFF 302273 PT<sup>*9</sup> | 2 | *Acer sp.* | AEAO 00000000 |
|  | N/A | Cit7<sup>*10</sup> | 2 | citrus leaf surface | AEAJ 00000000 |
|  | N/A | TLP2 | 2 | potato leaf surface | Gp0012374<sup>◇</sup> |
| Pph1448a | *phaseolicola*<sup>‡</sup> | 1448a<sup>*11</sup> | 3 | *Phaseolus vulgaris* | CP000058 |
| Pph1644r | *phaseolicola*<sup>‡</sup> | 1644r<sup>*12</sup> | 3 | *Vigna radiata* | AGAS 00000000 |
| PmaES4326 | *maculicola*<sup>¥</sup> | ES4326<sup>*13</sup> | 5 | *Raphanus sativus* | AEAK 00000000 |
| PsyUB246 | *syringae* | UB246<sup>*5</sup> | 13 | Fresh stream water | AVEQ 00000000 |

‡Pathovar is renamed as *Pseudomonas savastanoi*, not *P. syringae*.
¥ Strain is renamed as *Pseudomonas cannabina* pv. *asileensis*.
◇ JGI project id, not GenBank accession.
\*References for strain isolation information where available:
<sup>1</sup>(Cuppels 1986); <sup>2</sup>(Mukoo 1955); <sup>3</sup>(Baltrus et al. 2011); <sup>4</sup>(Morris et al. 2008); <sup>5</sup>(Morris et al. 2010); <sup>6</sup>(Loper and Lindow 1987); <sup>7</sup>(Mott et al. 2016); <sup>8</sup>(Berge et al. 2014); <sup>9</sup>(Sawada et al. 1999); <sup>10</sup>(Hirano and Upper 1990); <sup>11</sup>(Teverson 1991); <sup>12</sup>(Baltrus et al. 2012); <sup>13</sup>(Davis et al. 1991).

Pathogenicity assays utilizing artificial aphid diet (Prosser and Douglas 1991) were carried out as detailed in Smee et al. (2017), with no alterations. Further details are given in Supplementary Online Materials. Briefly, bacterial suspensions at an OD_600_ of 0.8 were added in a 1:5 ratio to artificial aphid diet with 10 mM MgCl_2_ used for the negative control treatment. Feeding sachets of diet (and bacteria) were inverted above 5-6 day old, age-controlled aphids singly placed in wells of a 96-well plate, to allow monitoring of individuals. Assay plates were kept at 20°C and every 24 hours the feeding sachet was replaced with another sachet of sterile diet only. At 72 hours, which has been previously shown to be indicative of virulence differences between strains (Smee et al. 2017), mortality was recorded. Assays were replicated a minimum of three times.

Assays utilizing artificial diet can be conducted in controlled environments without additional biotic or abiotic factors. However, to demonstrate that the mortality rates on artificial diet aren’t an artifact of keeping aphids on diet for extended periods, we infected 100 aphids on diet but moved them after 24-hours to fresh two-week-old *V. faba* plants, and left them for another 48-hours at 20°C. This was repeated twice for four strains, two that are considered highly pathogenic to aphids, and two that are less so (PsyB728a; Cit7; PtoDC3000; and Pph1448a) as well as a control group that had 10 mM MgCl_2_ buffer mixed with artificial diet instead of bacterial solution.

### 2.3 Epiphytic assays

To assess epiphytic ability, overnight bacterial cultures were resuspended to a final OD_600_ of 1.0 in approximately 20 mL of 10 mM MgCl_2_. This was then sprayed onto the upper and lower leaf surfaces of eight to ten leaf pairs of *V. faba* (three to four individual plants in one pot) using a fine mist spray bottle, until run-off. Once fully dried, plants were incubated at 22°C and 90 ± 5% humidity for 48 hours. From each pot of *V. faba*, ten 1.27 cm diameter discs were taken using a sterilized cork borer from across the leaf pairs for each of three replicates, and placed into 10 ml of 10 mM MgCl_2_ buffer, then sonicated for 10 minutes and vortexed to dislodge epiphytic bacteria. Three serial dilutions were plated from each replicate, which were averaged to give a CFU count per 10 ml for each strain. The entire process was replicated a minimum of three times for each strain (four replicates were achieved for two pathovars: PsyB728a and Psy1212). Only CFU counts from dilutions containing 5-50 countable colonies were included, to minimize errors.

### 2.4 Impact of aphid presence on plants

Eight strains varying in epiphytic ability and virulence to aphids were tested to determine if the presence of aphids on plants benefits epiphytic strains of *P. syringae.* These strains included four that were deemed to be good epiphytes and highly virulent to aphids (PsyB728a; PsyCC1458; Cit7; and TLP2), and four that are poorer epiphytes and less virulent to aphids (PtoDC3000; Ptt50252; PsyCC1543; and Pph1448a).

Overnight bacterial cultures were prepared to a final OD_600_ of 0.8 in approximately 40-50 ml of 10 mM MgCl_2_. Three two-week old *V. faba* plants, each with at least ten leaf pairs, were sprayed as before. After 48 hours, long enough for the epiphytic bacterial populations to establish well, 200 age-controlled 5-6 day old aphids were added to one of the three plants. The aphids were placed on the soil in a central location and the plant covered with a perforated 25.4 x 40.6 cm bread bag. A second plant was also covered with a bread bag, but without any aphids. The final plant was taken to be processed as before for a 48-hour baseline CFU count (per 10 ml). Finally, an unsprayed two-week old *V. faba* plant had 200 aphids added, a bread bag covering it, and it was placed with the two remaining plants as a control for aphid survival. Two days later, twenty aphids were sampled from each of the sprayed plants and crushed individually with a pestle in 100 μl 10 mM MgCl_2_ and then the entire suspension plated with sterile glass spreading beads on King’s B media with added rifampicin (50 ng/μl). Colonies were checked 2-3 days later, and an individual was categorized as infected if more than ten CFUs were present. Twenty aphids from the control plant were removed to ensure numbers were kept equal across all plants. The experiment ended after a total of seven days of which aphids were present on plants for five. The entire process was repeated three times for each strain. At the final time point we measured aphid survival on the control, unsprayed, plant as well as the bacterial plants, and again tested twenty aphids for infection, as before. We also took measurements of epiphytic CFU counts as before, using ten leaf discs in three replicates per plant.

Previous studies have shown that aphids can become infected from epiphytic bacteria on leaf surfaces (Stavrinides et al. 2009; Hendry et al. 2018). Consequently, we hypothesized that the mortality of aphids feeding on epiphytically infected plants should be similar to that which we see when feeding bacteria suspended in artificial diet to aphids, yet likely take longer to achieve as aphids would be less likely to become rapidly or heavily infected. To demonstrate this for one pathovar (PsyB728a), we moved the surviving adult aphids at the end of an experiment onto fresh two-week-old *V. faba* plants for another four days. Moving the aphids to fresh plants meant that it was easier to distinguish the surviving original adults from their offspring over a longer time frame. This was repeated three times.

### 2.5 Honeydew assays

At least one strain of *P. syringae* grows to high titers inside aphids and is secreted in aphids’ honeydew (PsyB728a: Stavrinides et al. 2009). We therefore investigated whether bacterial growth inside aphids leads to higher epiphytic populations and if honeydew is beneficial to epiphytic *P. syringae*. We used two of the strains that benefitted most from the presence of aphids: Cit7, which is highly virulent to aphids, and PtoDC3000, which is much less virulent.

Bacterial cultures at a final OD_600_ of 0.8 were sprayed onto *V. faba* leaves, as well as added to artificial diet at a ratio of 1:5 bacterial culture:diet. The experimental design, described in detail in the Supplementary Online Materials, allowed us to eliminate the effect of aphid feeding inducing plant-mediated effects, as the honeydew was able to fall onto leaves with epiphytic bacteria, but the aphids were unable to feed on the leaves (Figure S1). This was achieved by separating aphids feeding on artificial diet from sprayed leaves of *V. faba* with a piece of fine mesh. There were three treatments replicated three times within each experimental block, which was also repeated independently three separate times. The three treatments were as follows: the entire set up with just sterile diet in the feeding sachet and no aphids present; the same but with aphids present; and finally, both with aphids present and also with the bacterial culture added to the feeding sachet for the first 24-hours. All feeding sachets were refreshed after 24-hours, and then left for a further 48-hours before the leaves were processed for epiphytic growth as before. The dishes were kept at 22°C and 90 ± 5% humidity for the duration of the experiment, and under UV-blocking plastic for the first 24-hours whilst bacteria were present, as aphids avoid fluorescent strains of *P. syringae* and hence may feed less (Hendry et al. 2018). To further investigate whether strains of *P. syringae* can utilize aphid honeydew as a resource, we grew all eight strains from the previous section in artificial honeydew medium. This medium included diluted artificial aphid diet, so that the sucrose content was reduced to the equivalent of 2% w/v and it contained low levels of other nutrients found in honeydew (Leroy et al. 2011*b*), and then we added fructose at 2% w/v and glucose at 4% w/v as suggested by Sinka et al. (2009). We note that in nature honeydew composition will likely vary depending on factors related to both the aphid and host plant (Leroy et al. 2011*b*; Pringle et al. 2014). Our goal was not to achieve a full representation of pea aphid honeydew, but to recapitulate key nutrient and sugar factors. Overnight bacterial cultures were resuspended to a final OD_600_ of 0.8 and 20 μl added to artificial honeydew for a final volume of 180 μl and incubated at 22°C with constant shaking on a Bioscreen C^TM^ (Growth Curves USA) to generate growth curves over a period of 48 hours. To determine intrinsic growth rate of these strains, we repeated the growth assay above using just KB media with added rifampicin (50 ng/μl).

Additionally, in order to eliminate possible plant-mediated effects from the benefit strains gained during our original experiment on whole plants, we conducted the same experiment but instead of the addition of aphids in one treatment, we sprayed plants with artificial honeydew. The honeydew was sprayed from a distance of approximately 15 cm so that individual droplets were distributed across leaves and the plants were immediately placed back in high humidity. As before, these were sprayed with honeydew after an initial period of 48-hours and then left another five days before processing and counting CFUs to determine if there was any benefit. The two strains that benefitted most in the first experiment were again used for this assay: Cit7 and PtoDC3000. This assay was repeated three times.

### 2.6 Statistical Analyses

All statistical analyses were conducted in R version 3.5.2 (R Core Team 2018). In all analyses, experimental block was initially included as a fixed effect and removed if non-significant, unless otherwise stated. In addition, all models were checked for normality of residuals and heteroscedasticity and were fine unless otherwise stated. All epiphytic growth data (colony-forming units: CFUs) were log_10_-transformed for analyses. All posthoc tests were assessed using the R package ‘emmeans’ (Lenth 2019), which uses a Tukey adjustment for multiple comparisons and a significance level of 0.05.

To discern the benefit of aphid presence to epiphytic bacteria, two-tailed t-tests were conducted on each pair of ‘with aphids’ and ‘no aphids’ at the 5-day timepoint. For each strain, the epiphytic CFU value of the ‘no aphids’ treatment was then subtracted from the value of the ‘with aphids’ treatment to give a measure of benefit. Relative benefit for each strain was calculated by dividing benefit by the ‘no aphids’ CFU value. Where possible, data from individual experimental blocks were all used in analyses as there was much variation across blocks. However, for some Pearson’s correlations with other experimental data, mean values for each strain had to be used.

Differences in virulence to aphids, epiphytic ability, infection rates, survival on plants, honeydew deposited on leaves and bacterial growth in artificial honeydew medium were all analyzed using General Linear Models (GLMs) with Gaussian distributions. Response variables were either proportion of aphids dead or infected, epiphytic growth data, or OD_600_ values at the 48-hour time point for bacterial growth. Fixed explanatory effects were bacterial strain in most analyses, with time period as well for comparing infection frequencies. When comparing survival after nine days on plants, a Welch’s Two-Sample t-test was used due to unequal variances. To investigate honeydew effects, the two strains were analyzed separately as we were not interested in comparing across strains, only within treatments.

## 3 Results

### 3.1 Epiphytic ability correlates with virulence to aphids

Some of the data presented here have been previously published by Smee *et al.* (2017). This includes both pathogenicity to *A. pisum* and epiphytic assays for nine strains (Table 1, highlighted in grey: PsyB728a; PtoDC3000; Pja301072; Ptt50252; Ppi1704B; Pac302273; Cit7; TLP2; and Pph1448a). For the current study, this was expanded to include another 12 strains, and replicates for epiphytic assays were increased from two to a minimum of three.

Across the 21 strains of *P. syringae* there was much variation in both their epiphytic ability on *V. faba* and their pathogenicity to aphids (Figure 1: GLM: *F*_20,171_ = 22.56, *p* < 0.001; GLM: *F*_20,72_ = 5.31, *p* < 0.001, respectively). Some of the best epiphytes were capable of colonizing *V. faba* leaves at over 1 × 10^7^ CFUs per 25.8 cm^2^ area of leaf (the upper and lower surfaces of 10 leaf discs each with 1.27 cm diameter). Three of these strains (PsyB48, PsyUSA011 and Cit7) were also amongst the most pathogenic to aphids, causing over 74% death after 72 hours post-infection by artificial diet. Similarly, some of the poorest epiphytes were far less virulent to aphids, indicating that the two traits may be linked and there may be a common factor important for success or failure in both environments (Figure 2; Pearson’s product-moment correlation: *t* = 4.21, d.f. = 19, *p* < 0.001, *R* = 0.69). Note that the correlation is not due to aphids becoming infected more by the better epiphytes, since the mortality data is from controlled conditions in artificial diet, not from on plants. Also, the epiphytic ability of these strains may differ in nature, as our assays started with high founding populations of epiphytic bacteria and were kept at high humidity and ideal conditions for growth of *P. syringae*. Hence, our results may reflect maximum densities achievable on fava bean plants.

**Figure 1.**
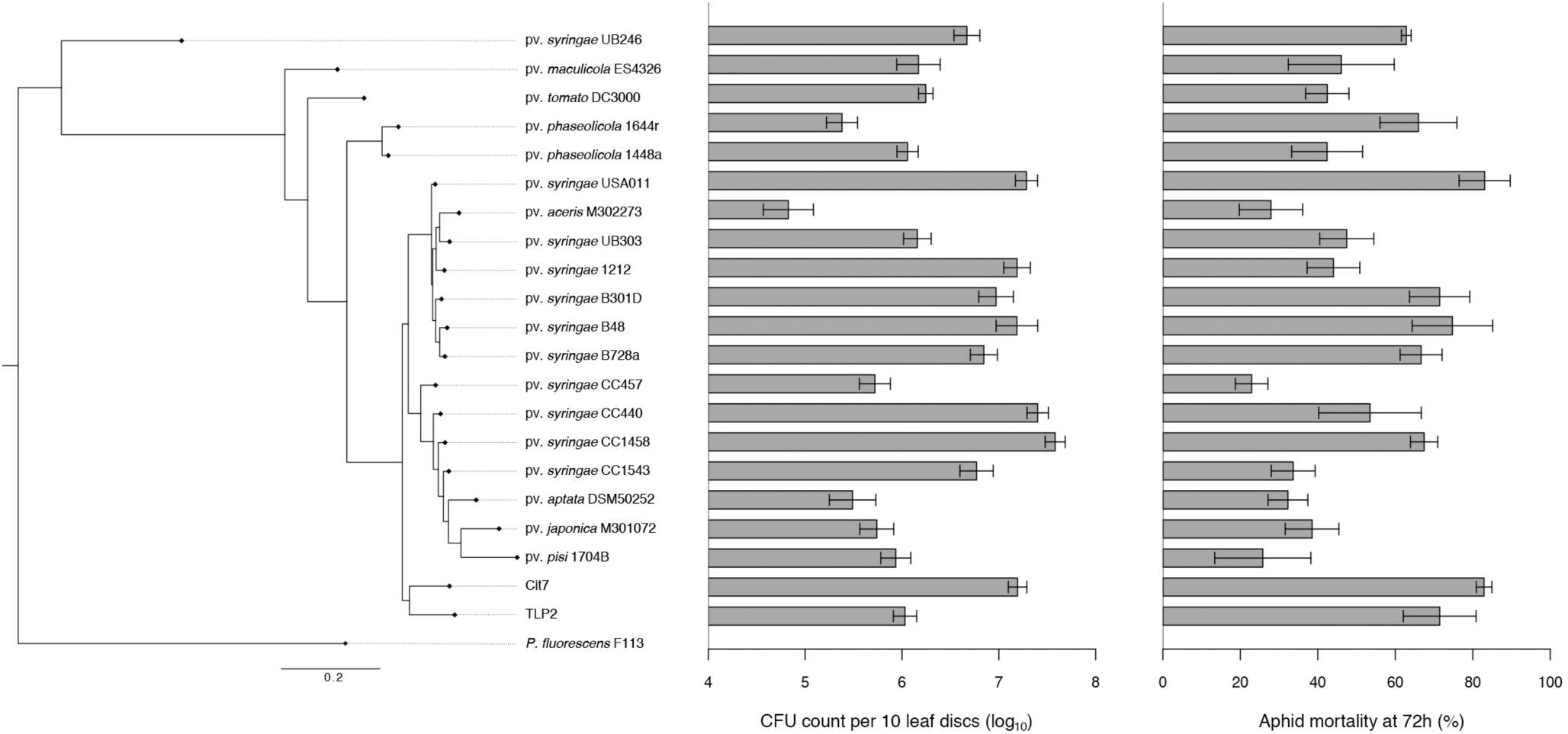
A comparison of epiphytic ability and virulence to aphids across 21 strains of *P. syringae.* The phylogenetic tree was constructed in PhyloPhlAn (Segata et al., 2013) using 382 of the most conserved protein sequences across the 21 genomes, and rooted using the representative genome of *Pseudomonas fluorescens* F113 from GenBank (accession number NC_016830). All other protein sequences were also obtained from GenBank (see Table 1). The epiphytic ability of the 21 strains is measured as log_10_ CFU counts per 10 leaf discs sampled, and the *y*-axis starts at the minimum CFU count we might see in our dilutions, given accuracy cut-offs of 5-50 colony counts per 5 μl droplet, taken from an initial 10 ml sample. The virulence of each strain to aphids is represented by aphid mortality at 72 hours post-infection in artificial diet. Mean values are plotted ± standard error.

**Figure 2.**
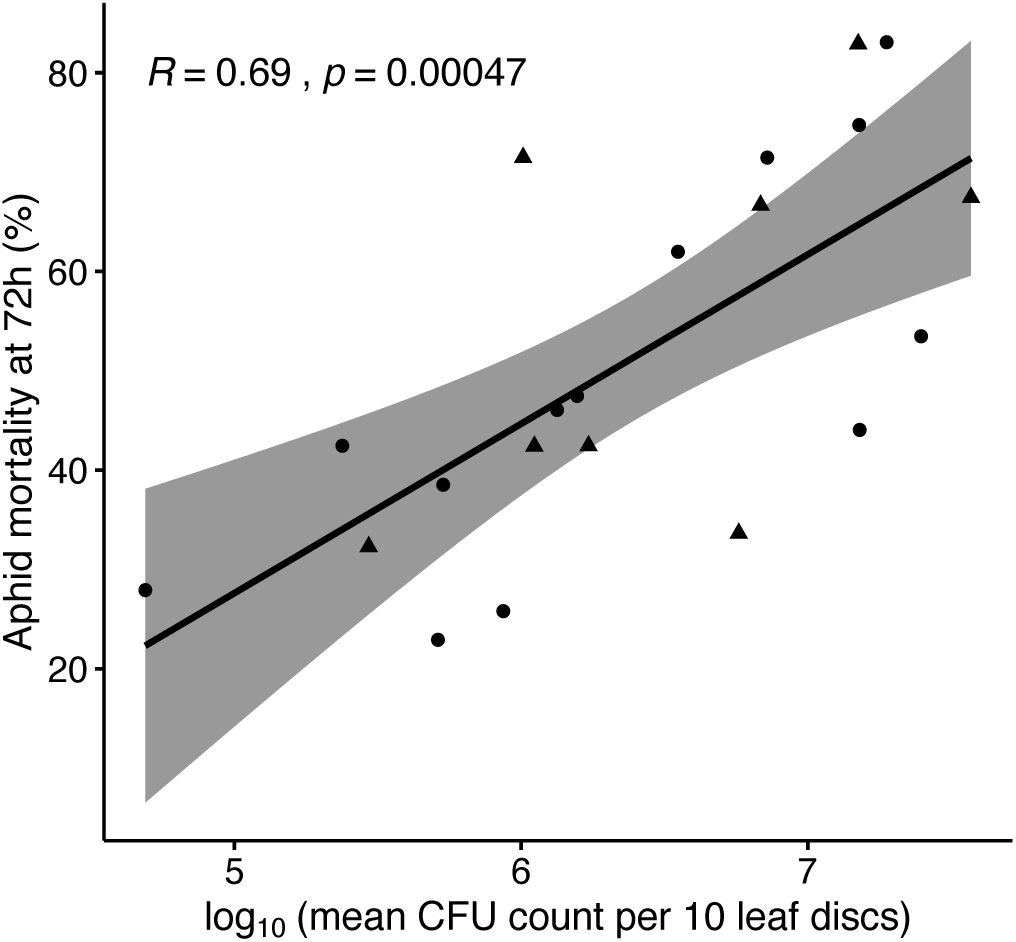
Correlation between the epiphytic ability of each of the 21 *P. syringae* strains and their virulence to aphids after 72 hours. Pearson’s correlation coefficient (*R*) is presented, and the grey area denotes 95% confidence intervals. Those strains used in further experiments are shown as triangular points.

### 3.2 Aphid presence can benefit epiphytic bacteria

To determine if the correlation between epiphytic ability and virulence could be adaptive, we investigated whether strains of *P. syringae* growing epiphytically actually benefitted from the presence of aphids on plants. If using aphids as hosts is a viable ecological and evolutionary route for the bacteria, we would expect that there would be some benefit gained from the two approaches. Better epiphytes kill their hosts more quickly, so perhaps they would more quickly return to their preferred niche of leaf surfaces. Poorer epiphytes stay in the living host for longer, which may increase their epiphytic titers by dispersal via the honeydew secreted by the hosts.

We selected eight strains from various points along the correlation in Figure 2, denoted by triangular points on the plot. When comparing CFU counts from *V. faba* plants either with or without aphids present for five days, several of the strains benefitted and demonstrated significant increases in bacterial densities (Figure 3A). However, when comparing this benefit (change in CFU counts) to the virulence of each strain, there was no correlation (Pearson’s product-moment correlation: *t* = 0.51, d.f. = 6, *p* = 0.63, *R* = 0.21). There was also no correlation with the epiphytic ability of each strain, defined using epiphytic data in Figure 1 (Pearson’s product-moment correlation: *t* = 0.24, d.f. = 6, *p* = 0.82, *R* = 0.10). Using the relative benefit gained by each strain (benefit divided by epiphytic CFU counts without aphids present) did not change these results.

**Figure 3.**
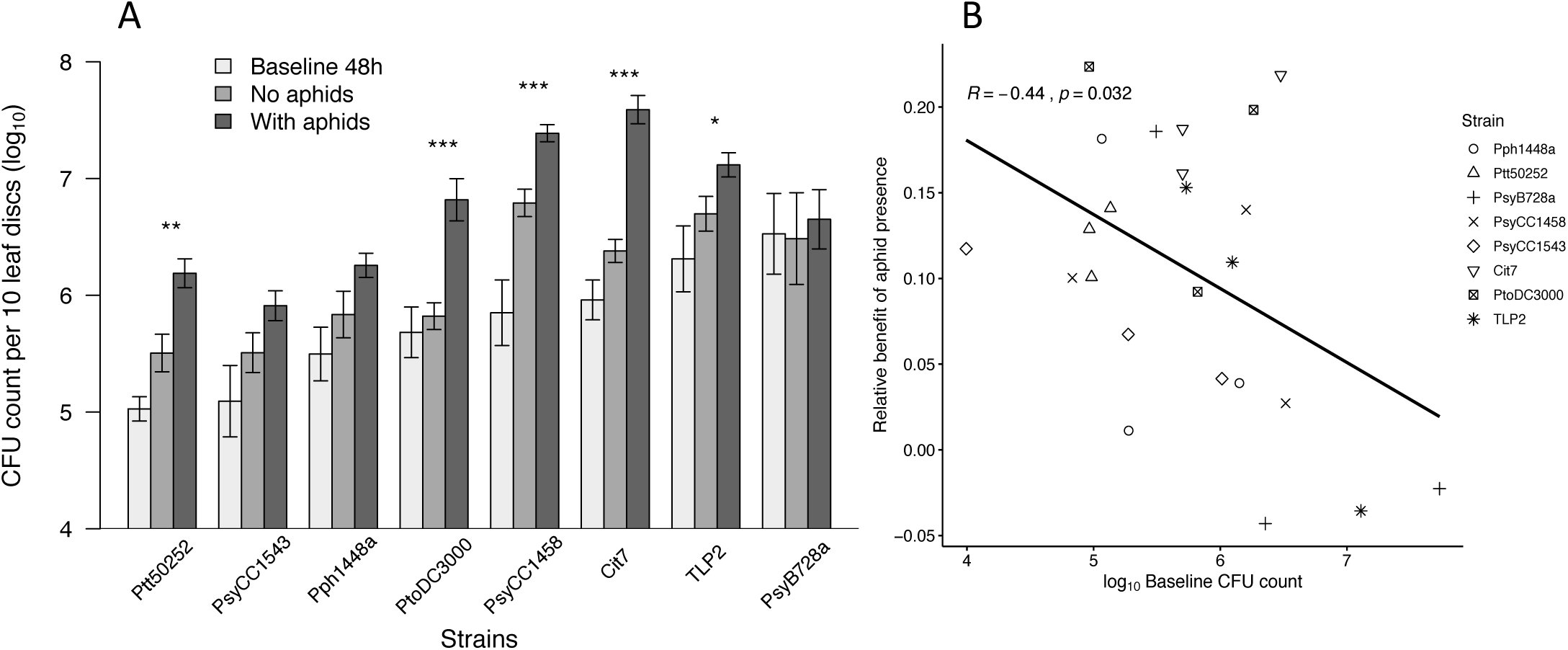
Benefit of aphid presence. (A) Epiphytic CFU counts for each strain sprayed on to fava beans. Baseline values are after 48-hours, then aphids were added and five days later CFU counts were taken from plants with and without aphids. Strains are ordered by 48-hour baseline values, smallest first. Mean values are plotted ± standard error. Significances are given between treatments of ‘no aphids’ and ‘with aphids’ at levels of: * *p* < 0.05; ** *p* < 0.01; *** *p* < 0.001. (B) The relative benefit of aphid presence after five days, irrespective of strain, is negatively related to baseline epiphytic CFU counts at 48-hours (LM: F_1,22_ = 5.28, *p* = 0.03, intercept = 0.35, slope = −0.04, *R* = −0.44).

However, the baseline epiphytic growth at the 48-hour timepoint was a very strong predictor of the relative benefit gained after five days, when using data from all experimental blocks (Figure 3B; Linear model: *F*_1,22_ = 5.28, *p* = 0.03). Interestingly, there was a negative relationship indicating that replicates where bacteria benefitted the most from the presence of aphids had lower epiphytic growth earlier in the experiment, and vice versa.

It is also possible that the ability to infect aphids impacts success for epiphytic bacteria such as *P. syringae*. If some strains are less able to infect hosts from the phyllosphere, then they have less chance of growing within the host at all and being dispersed via honeydew. To confirm that aphids actually become infected from epiphytic populations of *P. syringae* we sampled aphids after being on infected plants for 48-hours and also after five days (Figure 4A). All eight strains were able to infect aphids, and infection rates on plants were much higher after five days than 48-hours (GLM: *F*_1,39_ = 32.29, *p* < 0.001). There was a large range in mean infectivity levels after five days, from 23% for strain Ptt50252 to 100% for strain PsyCC1458, which is tightly linked to the mean epiphytic density of bacteria at the same time point (Figure 4B; Pearson’s product-moment correlation: *t* = 4.45, d.f. = 6, *p* = 0.004, *R* = 0.88), but not to the benefit (change in CFU counts) of aphid presence (Pearson’s product-moment correlation: *t* = 0.43, d.f. = 22, *p* = 0.67, *R* = 0.09).

**Figure 4.**
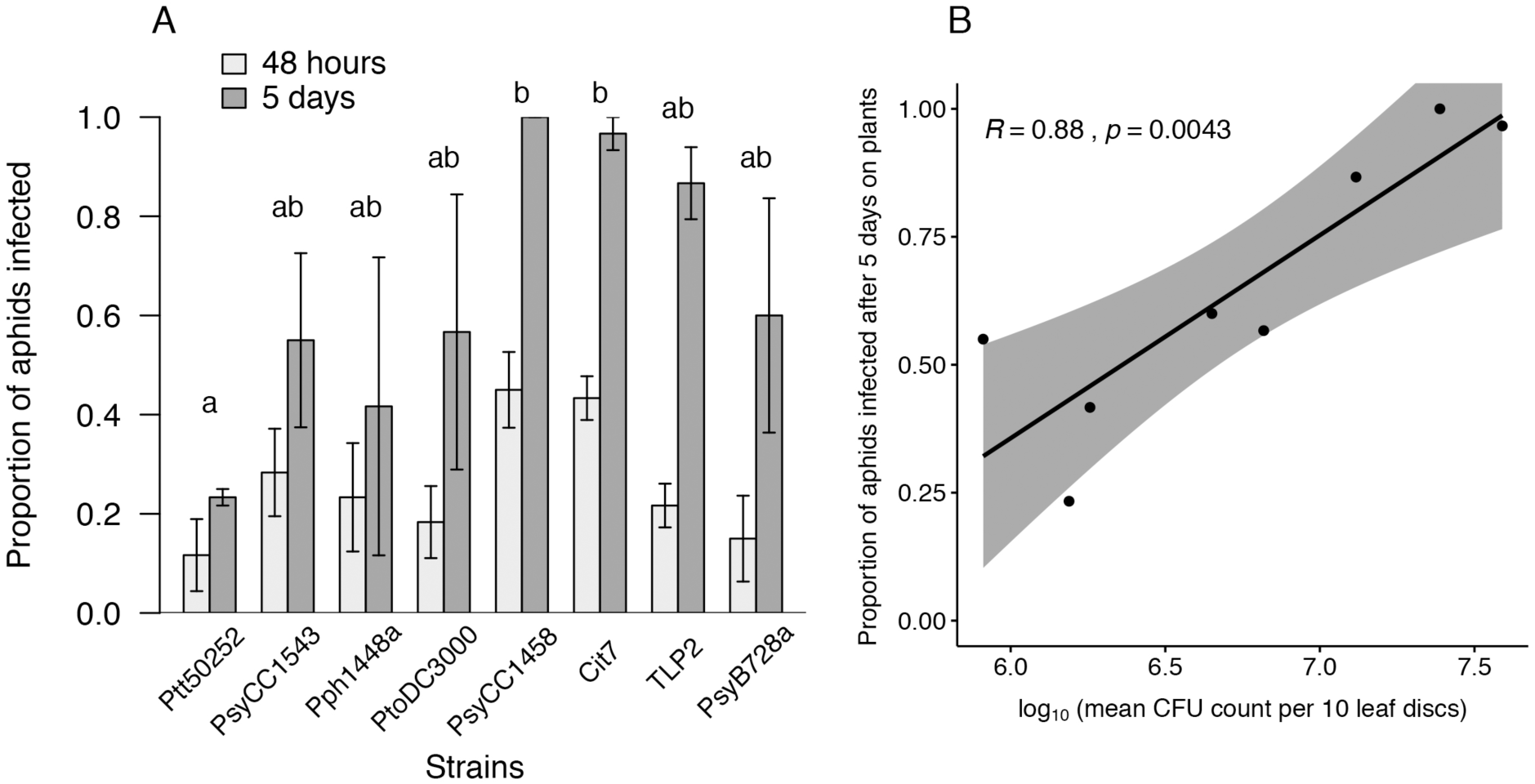
Aphid infection rates on plants. (A) After 48-hours and 5 days on plants sprayed with epiphytic bacteria. Mean values are plotted ± standard error. Posthoc lettering denotes overall differences between strains, irrespective of time, at p = 0.05 level. (B) Infection rates at 5 days on plants were highly correlated with the epiphytic bacterial population on the leaves at that time (data from ‘with aphids’ treatment from Figure 3; Pearson’s product-moment correlation: *t* = 4.45, d.f. = 6, *p* = 0.004, *R* = 0.88).

In previous experiments, epiphytic ability was connected with virulence on artificial diet, not virulence directly from epiphytic populations of bacteria. Hence, we investigated whether aphid mortality on plants was similar to that on artificial diet, and therefore if our expectations would be the same. After five days on infected plants, there was no difference in aphid mortality across the eight strains being tested, or when compared to a control treatment that had only buffer sprayed onto the leaves (Figure S2A; GLM: *F*_8,28_ = 0.52, *p* = 0.84). However, if aphids were infected on artificial diet initially, then placed onto *V. faba* plants for 72 hours, the mortality rates varied among strains and were significantly different to the control treatment, and closer to what we would expect from our previous data (Figure S2B; GLM: *F*_4,5_ = 33.18, *p* < 0.001). This confirms that it is not the feeding on artificial diet per se that promotes quicker death, but perhaps the uptake of more bacterial cells. Consequently, as the dosage of epiphytic bacteria ingested by aphids is likely much lower on plants than when fed artificial diet, we expected the effect to take longer when on plants. For one strain, PsyB728a, we extended the experiment on plants only and found that during an extra four days, to make nine days total on plants, a significant difference did emerge between aphid mortality on control plants and infected plants (Figure S2C; Welch Two Sample t-test: t = −4.83, df = 2.26, p = 0.03). We also observed transfer of bacteria onto the fresh *V. faba* plants from the aphids initially fed bacterial solutions via artificial diet.

### 3.3 Honeydew may provide a valuable resource

We hypothesized that bacteria could benefit both from growth inside aphids and also from honeydew, so we sought to determine the relative benefit of growth inside aphids versus increased epiphytic growth from the addition of honeydew to leaf surfaces by experimentally separating these components. We would expect that with the addition of bacteria growing inside the aphid host and being excreted in honeydew would amplify the populations of bacteria on the leaf surface. Alternatively, bacteria already on a leaf might benefit from additional nutrients deposited on the leaf in honeydew and grow to higher titers.

Two strains were chosen for this assay, PtoDC3000 and Cit7, as both had shown that they benefitted from the presence of aphids on plants (Figure 3) but differed in their virulence to aphids and their epiphytic ability (Figure 1). CFU values of both strains varied over experimental blocks (PtoDC3000: GLM: *F*_2,49_ = 7.59, *p* = 0.001; Cit7: GLM: *F*_2,49_ = 3.65, *p* = 0.03), but there was no significant interaction, so the pattern reported did not change. For both strains, the treatment that had no aphids present repeatedly gave the lowest CFU counts, compared to treatments with aphids, whether or not they were also feeding on bacteria (Figure 5A: PtoDC3000: GLM: *F*_2,49_ = 8.92, *p* < 0.001; Cit7: GLM: *F*_2,49_ = 4.95, *p* = 0.01). Therefore, at least for these two strains, the presence of honeydew appeared to increase population densities, but additional growth of bacteria inside the aphids did not contribute to the increased titers of bacteria on leaves.

**Figure 5.**
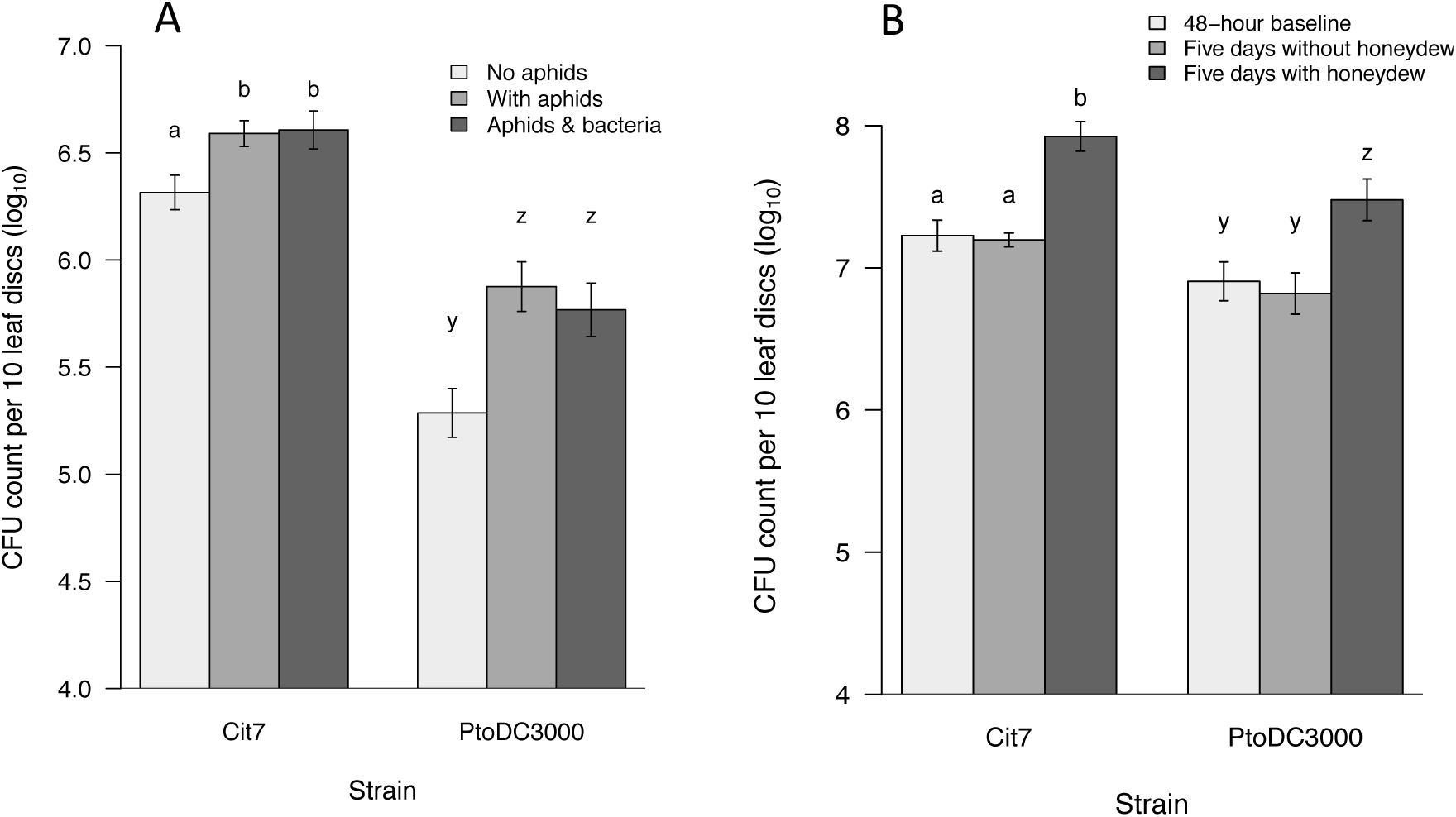
The effect of honeydew on leaves. (A) Epiphytic CFU counts from leaves sprayed with bacteria and then with either no aphids added, or aphids feeding on artificial diet suspended above the leaves, with or without added bacterial suspensions. The presence of aphid honeydew on leaves increases bacterial CFU load, regardless of whether or not the aphids had ingested bacteria. (B) Epiphytic CFU counts from whole plants 48-hours after being sprayed with bacteria, and then after another five days with or without artificial honeydew sprayed onto leaves. Even without the presence of aphids feeding, honeydew is beneficial to these epiphytic strains of *P. syringae*. Mean values are plotted ± standard error, and lettering indicates results of post-hoc tests, with strains analyzed separately.

We further demonstrate that it is honeydew that benefits the epiphytic bacteria, and not plant-mediated effects of aphids feeding on plants, by excluding aphids completely from our assays on whole plants. Both Cit7 and PtoDC3000 showed a significant increase in population densities when plants were sprayed with artificial honeydew and left for five days, compared to unsprayed control plants which did not differ from baseline values after 48-hours (Figure 5B: PtoDC3000: GLM: *F*_2,22_ = 10.48, *p* < 0.001; Cit7: GLM: *F*_2,22_ = 20.55, *p* < 0.001). Across experimental blocks there was some variation in PtoDC3000 densities, but not Cit7 (PtoDC3000: GLM: *F*_2,22_ = 8.98, *p* = 0.001; Cit7: GLM: *F*_2,22_ = 1.09, *p* = 0.36).

We then grew all eight strains in artificial honeydew, to confirm whether the ability to utilize honeydew as a resource leads to epiphytic benefit for particular strains. All eight strains grew well in the artificial honeydew medium. From a starting OD_600_ of 0.1 ± 0.01, strain PsyCC1458 grew the most, up to an OD_600_ of 1.73 in 48 hours, and strain PsyCC1543 grew the least, to an OD_600_ of 1.17 (Figure S3A). The eight strains showed a significant amount of variation in their ability to grow in the honeydew medium at the final time point (GLM: *F*_7,72_ = 219.77, *p* < 0.001), but also showed no correlation with the benefit gained by each strain in Figure 3 (Pearson’s product-moment correlation: *t* = −0.18, d.f. = 6, *p* = 0.87, *R* = −0.07). These growth patterns do not reflect the intrinsic ability of each strain to grow, as there is no correlation with their growth patterns in KB medium (Figure S3B).

## 4 Discussion

As perhaps the best-studied plant pathogen to date, and an influential model system for studying plant-microbe interactions and the evolution of virulence, *P. syringae* provides a powerful tool for exploring more complex, tripartite interactions in the phyllosphere. More recent genomic tools have revealed the impressive diversity of this bacterium in nature (Baltrus et al. 2011; Berge et al. 2014; Thakur et al. 2016). However, much of the literature has focused on revealing variation in pathogenicity traits associated with plants (Hwang et al. 2005; Lindeberg et al. 2012). We show that in addition to this, across 21 strains of *P. syringae* that vary in plant virulence, epiphytic growth on fava beans is also highly variable and strongly correlated with virulence to aphids, building on a previous study by Smee et al (2017). This would suggest that the two traits are somehow linked and that those strains most often encountered on plants are likely to be entomopathogenic.

Out of the eight strains chosen to further investigate this relationship between epiphytic ability and virulence to aphids, five were shown to significantly benefit from the presence of aphids on plants. If gaining a benefit from the presence of aphids is adaptive, we hypothesized that there may be a trade-off between virulence and dispersal, in that less virulent strains remain in the aphid host for longer, allowing more dispersal, and in which case those strains might benefit more from the presence of aphids. We found that this was indeed the case for these *P. syringae* strains as a whole, although irrespective of specific strain. Epiphytic growth at 48-hours was found to significantly predict the relative benefit aphids provided to bacteria after being on plants for five days, with poorer epiphytes benefitting more. As there was much variation across experimental blocks within single strains this effect was lost if only using the mean of each strain, suggesting that the extent to which strains benefit is likely context dependent. There is often a lot of variability in how cells react to different environmental stimuli, so drawing overarching conclusions about each strain at this point is difficult.

Low virulence to aphids may have evolved in poorer epiphytes that benefit more from keeping aphids alive for longer. Better epiphytes that don’t benefit from the presence of aphids can consequently evolve high virulence. When correlating the mean benefit gained by each strain against the mean virulence to aphids, or our initial measure of epiphytic ability (sprayed at an OD_600_ of 1.0), there were no significant relationships, although this could again be due to losing variation through the use of mean data points for each strain. Short-term mortality of aphids on plants with epiphytic bacteria was found to be less extreme than when aphids were infected on artificial diet. This was somewhat unsurprising, assuming aphids ingest far more bacterial cells when feeding on artificial diet than when feeding on infected plants. However, over longer time periods, the relative differences in virulence between the strains is likely to be the same as on artificial diet as infection rates were shown to be highly correlated with epiphytic growth at five days, and once infected aphids cannot recover. We also show, for one pathovar PsyB728a, that over another few days mortality is closer to the level seen on artificial diet, and significantly different to controls. It is possible that we would see differences in how strains benefit over longer time scales when aphid death would vary more between bacterial strain treatments. Alternatively, it is possible that the relationship between epiphytic ability and virulence observed here occurs because whatever trait causes a strain to grow well on a leaf surface is also the same trait that causes them to grow well inside an aphid and causes death to the aphid host. Given the lack of correlation between epiphytic growth or virulence and benefit from aphids found here, we conclude that there is no adaptive reason for at least some *P. syringae* strains to use aphids as alternative hosts.

In further support of the idea that *P. syringae* does not benefit from using aphids as an alternative host, we found that there was no additional impact on epiphytic cell titers of bacterial growth inside the aphids when honeydew was deposited on leaves, for the two strains tested. If the growth inside an aphid is not the source of benefit to epiphytic *P. syringae*, then ultimately it is only the bacterial cells already on the leaf surface that benefit from aphids. It is possible that strains other than Cit7 and PtoDC3000 might show different patterns, or that with different experimental conditions we would see different effects. However, we did find support to conclude that some strains benefit from aphid presence due to honeydew excretion. It could be that particular strains of *P. syringae* have evolved mainly to utilize aphids not as hosts, but instead the sugar-rich honeydew waste secreted by aphids onto leaves, in order to boost bacterial densities on leaves. Our data indicate that strains of *P. syringae* differ in their ability to use honeydew as a growth medium, and that across the eight strains tested this was uncorrelated and therefore unpredictable from their growth in KB media. Hence, strains are differentially adapted to utilize this potentially abundant resource. However, their growth in the honeydew medium was unrelated to the benefit of aphid presence. It could be that our artificial honeydew was not realistic enough, or that as honeydew on leaf surfaces dries and becomes more concentrated, then different bacteria are better or worse equipped to utilize it. Honeydew is carbon-rich, and hence a potential energy source for many organisms in the phyllosphere, but the composition differs depending on both the species of aphid and also the host plant they feed on, which has further knock-on effects to other organisms such as ants that feed on honeydew (Leroy et al. 2011*b*; Pringle et al. 2014). *Cinara* species of aphid also spread honeydew which has been shown to encourage growth of bacteria, yeasts and filamentous fungi (Stadler and Müller 1996). If honeydew is important in structuring communities on both the leaf surface, but also the surrounding habitat (Stadler and Müller 1996; Milcu et al. 2015), then aphids are surely engineers of both the phyllosphere microbiome, and also the wider ecosystem.

Most studies to date that illustrate an effect of honeydew, or herbivores in general, on bacterial populations in the phyllosphere have not managed to remove the potential plant-mediated effects of herbivore presence (Stadler and Müller 1996, 2000). By restricting aphids from feeding on leaves, but allowing honeydew still to be deposited or by spraying artificial honeydew on to plants, we show that the significant increase in growth of epiphytic *P. syringae* is not due to plant-mediated effects caused by herbivore feeding (Humphrey and Whiteman 2019). Despite this, there may have been differential plant-mediated effects acting in response to different strains of *P. syringae* or even the presence of honeydew on the leaves in experiments using aphids. Plant responses to bacteria are variable, and in nature may make plants more susceptible to herbivores due to crosstalk between JA and SA (Humphrey et al. 2014), which in turn may also positively affect the abundance and diversity of bacteria on some plants or trees (Stadler and Müller 2000; Müller et al. 2003). However, here we show that aphids can interact directly with epiphytic bacteria to influence their population densities. We further speculate that in natural, diverse communities differences in the bacterial ability to utilize honeydew would also impact competition between bacterial strains and species to influence community composition.

In conclusion, our results support the hypotheses (i) that there is a link between *P. syringae* virulence to the pea aphid, *A. pisum*, and epiphytic growth ability across a diverse set of strains, (ii) there is a benefit of aphid presence to some epiphytic bacterial populations, (iii) this benefit is not due to plant-mediated effects, nor bacterial growth within aphid hosts, but likely an interaction between poorer epiphytes and the excretion of honeydew onto leaf surfaces. Success in the phyllosphere is crucial for many *P. syringae* pathovars and environmental strains. The ability to successfully utilize ephemeral resources such as carbon-rich honeydew secretions from aphids could provide a competitive advantage to epiphytic bacteria, leading to higher densities on leaf surfaces, greater opportunity to infect other hosts, and even further reaching and less explored consequences, such as attracting a plethora of other vectors or hosts (Farré-Armengol and Junker 2019).

