## Supplemental Information for "Insects as phyllosphere microbiome engineers: effects of aphids on a plant pathogen"

### Additional Materials and Methods

*In vitro pathogenicity assays*

For each aphid assay, bacterial treatments were prepared by growing an overnight culture that was then pelleted, washed in 10 mM $MgCl_2$ and then pelleted again and resuspended in 10 mM $MgCl_2$ to an $OD_{600}$ of 0.8. Bacterial suspensions were added in a 1:5 ratio to artificial aphid diet with 10 mM $MgCl_2$ used for the negative control treatment. A 96-well plate was filled with diet, 200 µl per well, and then covered by parafilm to make a feeding sachet. In each well of another 96-well plate, a single age-controlled aphid (5-6 days old, approximately third instar) was placed and the feeding sachet plate was then inverted above them to allow oral exposure to the bacteria and diet solution, whilst enabling us to record individuals separately throughout the assay. Assay plates were kept at 20°C and after 24 hours the feeding sachet was replaced with another sachet of sterile diet only. Each strain persisted well in the artificial diet during the 24 hours period and there was no indication from dilutions and CFU counts that any of the strains performed better or worse than others in the artificial diet. The diet was refreshed again after another 24 and 48 hours. Mortality at 72 hours has been previously shown to be indicative of virulence differences between strains (Smee et al., 2017), so data at this time point was used in analyses. An aphid was assumed dead if it had turned brown or was at the bottom of the well (not feeding) and did not move when agitated. Assays were replicated a minimum of three times.

*Honeydew assays*

Bacterial cultures at a final $OD_{600}$ of 0.8 were sprayed onto *V. faba* plants for a total of 18 leaf pairs, as well as added to artificial diet at a ratio of 1:5 bacterial culture:diet. The final experimental set-up (Figure S1) consisted of an upturned plastic tip box lid, 11.4 x 7.6 x 2.5 cm, with two leaf pairs sprayed with epiphytic bacteria placed inside. The leaf pairs had their petioles placed in a small amount of 2% agar at opposite corners of the dish, enabling most of the surface of the dish to be covered by leaf. A rectangular piece of mesh was then loosely placed over the dish and secured with an elastic band. For those treatments with aphids we then placed 150-180, depending on availability that day, age-controlled aphids on top of the gauze. The central 32 wells (8 x 4) of a 96-well plate were filled with 200 µl of artificial diet (with or without bacteria), covered in parafilm and inverted above the aphids as a feeding sachet. Parafilm was used to secure the edges and keep the feeding sachet attached to the leaf dish. This experimental design allowed us to eliminate the effect of aphid feeding inducing plant-mediated effects, as the honeydew was able to fall onto leaves with epiphytic bacteria, but the aphids were unable to feed on the leaves.

There were three treatments replicated three times within each experimental block, which was also repeated independently three separate times, giving a total of nine dishes in each block and nine mean data points overall for each treatment.

The three treatments were as follows:

1) the entire set up with just sterile diet in the feeding sachet and no aphids present;
2) the same but with aphids present;
3) both with aphids present and also with the bacterial culture added to the feeding sachet for the first 24-hours.

All feeding sachets were refreshed after 24-hours, and then left for a further 48-hours before the leaves were processed for epiphytic growth as before, except only two replicates of ten leaf discs could be taken due to there only being two leaf pairs in each treatment. The dishes were kept at 22°C and 90 ± 5% humidity for the duration of the experiment, and under UV-blocking plastic for the first 24-hours whilst bacteria were present, as aphids avoid fluorescent strains of *P. syringae* and hence may feed less (Hendry et al., 2018).

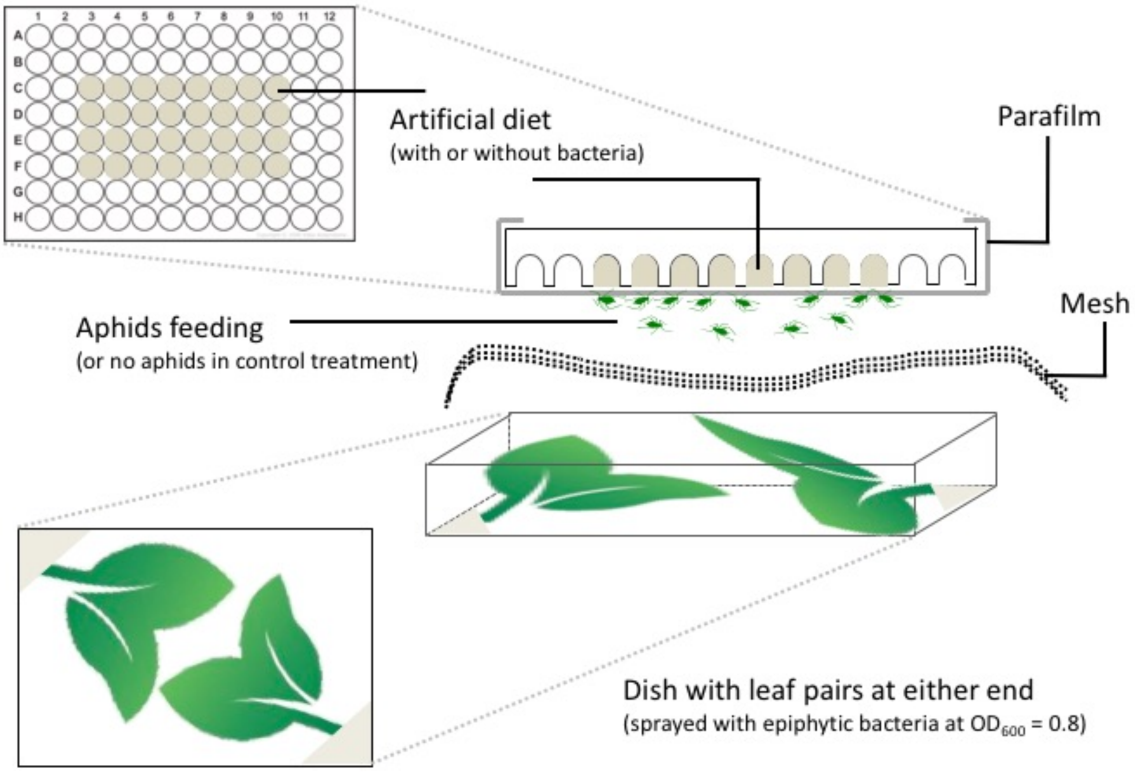


**Figure S1** – Experimental set-up for honeydew assays, to exclude aphid induced plant-mediated effects.

### Additional figures

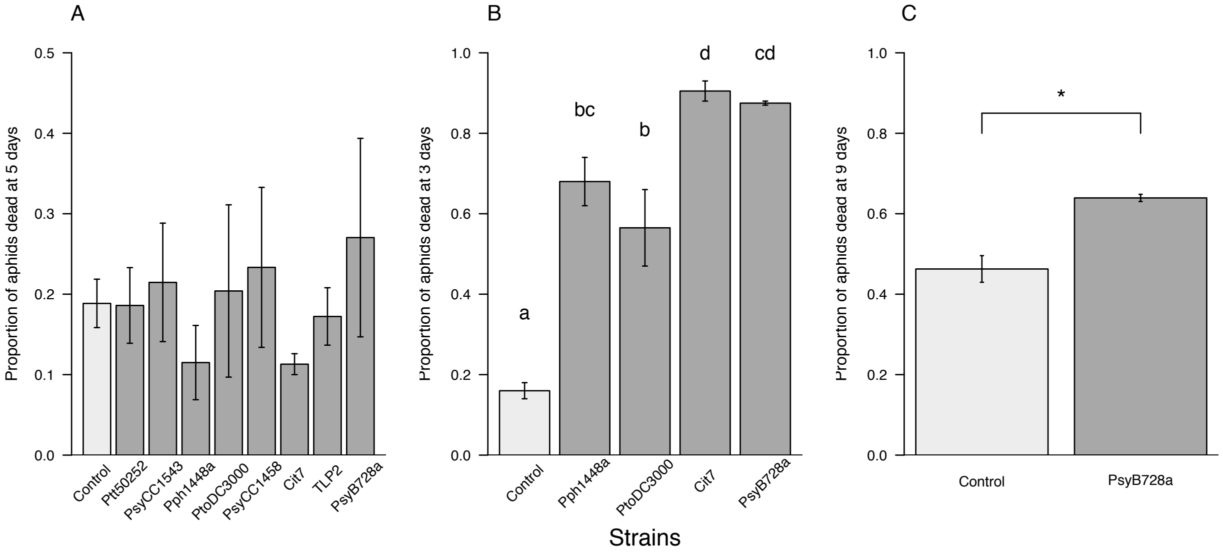


**Figure S2** – Aphid mortality (A) after five days on plants sprayed with epiphytic bacteria. (B) Aphids infected via artificial diet for 24-hours, then transferred to *V. faba* plants for another 48-hours exhibit similar levels of death than if left on artificial diet alone after infection. Letters denote results of posthoc tests. (C) Longevity of aphids infected on plants and left for an extra four days to make a total of nine days, for just strain PsyB728a, which differs significantly from the controls at the level of: * $p < 0.05$. Mean values are plotted ± standard error.

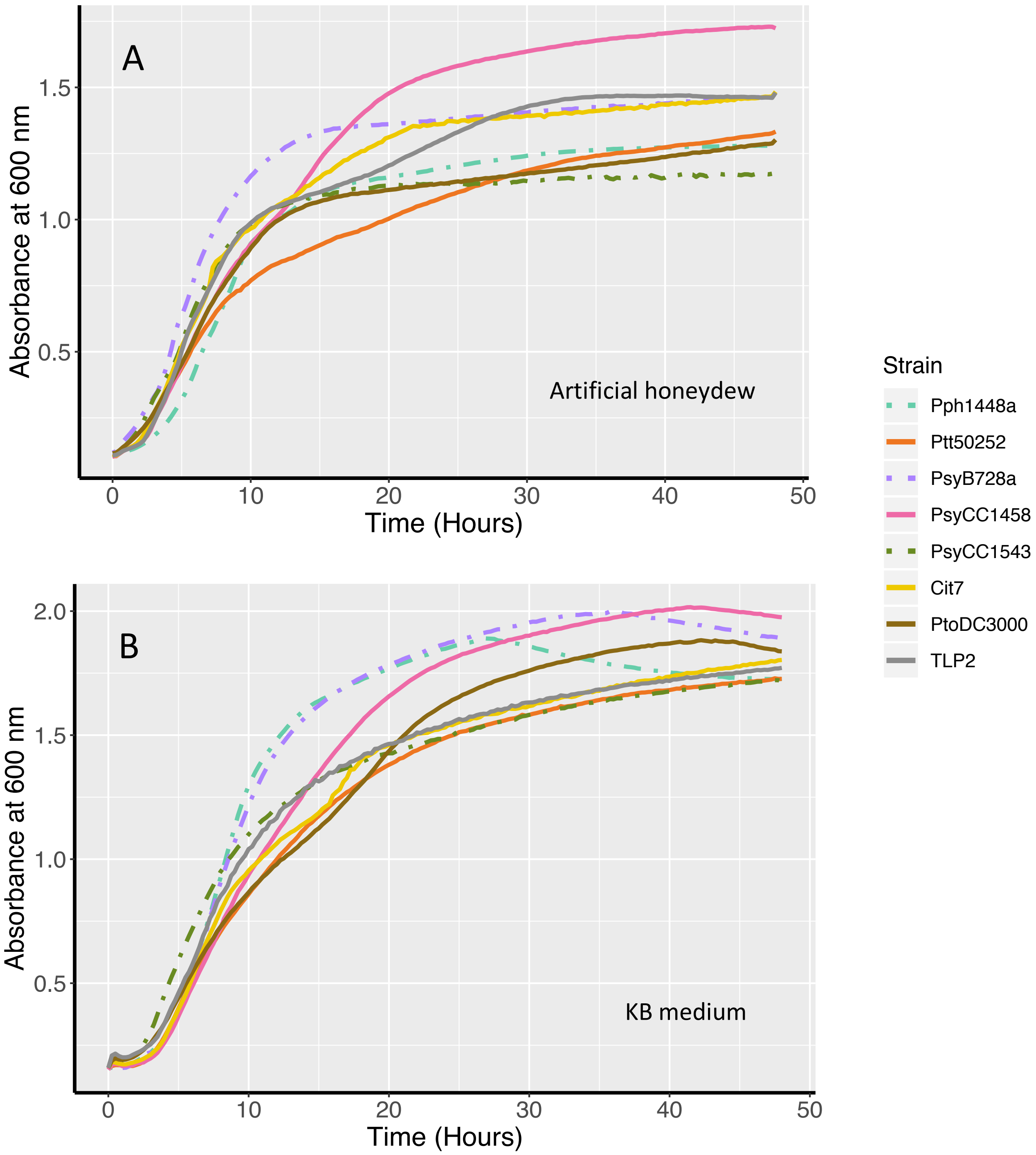


**Figure S3** – Growth of eight strains of *P. syringae* in artificial honeydew media (A), and in KB + rifampicin media (B), over 48 hours. Those strains that were shown to benefit from the presence of aphids are plotted in solid lines, and those that do not benefit are plotted in dashed lines.
